# Behavioral and genetic correlates of heterogeneity in learning performance in individual honeybees, *Apis mellifera*

**DOI:** 10.1101/2023.10.23.563551

**Authors:** Neloy Kumar Chakroborty, Gérard Leboulle, Ralf Einspanier, Randolf Menzel

**Affiliations:** Institute Biology, Neurobiology, Freie Universität Berlin, Königin Luisestr. 1-3, Berlin, Germany; Institute of Veterinary Biochemistry, Department of Veterinary Medicine, Freie Universität Berlin, Oertzenweg 19b, Berlin, Germany

## Abstract

Learning an olfactory discrimination task leads to heterogeneous results in honeybees with bees performing very well and others at low rates. Here we investigated this behavioral heterogeneity and asked whether it is associated with particular gene expression patterns. Bees were individually conditioned in a sequential conditioning protocol involving several phases of olfactory learning and retention tests. The rate of CS+ odor learning was found to correlate highest with a cumulative performance score extracted from all learning and retention tests. The cumulative score was used to sort the tested bees into high and low performers. Microarray analysis of gene expression in the mushroom body area of the brains of these bees disclosed a list of genes, which were differentially expressed between the high and low performers. These candidate genes are implicated in diverse biological functions, such as neurotransmission, memory formation, cargo trafficking and development.

## Introduction

Central components of animal cognition are their ability to adapt to new environmental conditions, store successful changes of their behavior caused by new learning, and use the relevant memories for further behavioral control. In honeybees, a classical model of learning and memory studies, a large body of data document how cognitive functions are essential for their foraging success and survival of their colonies [1] [2]. Evolutionary selections for the improvements of cognitive abilities work on the individual level and are based on genetic variations that are favorable for behavioral traits related to sensory processing, learning and memory formation [3] [4]. The search for such cognitive adaptations can be performed both on a population level and on an individual level. However, little efforts have been concentrated on the study of the causes and lifestyle-related effects of individual variations in cognitive abilities and styles within single species [5] except for humans [6] [7] [8]. Studies across different vertebrate species although recognized the consistency of inter-individual variability in cognitive capacities ([9] [10] [11]), albeit tracing invertebrate cognition through the lens of individual behavioral variability is a much less explored.

Selection for behavioral traits usually involve many animals. Studies on learning and memory in *Drosophila* have led to the discovery of multiple genes related to learning and memory and groups of flies belonging to the same line can be tested sequentially for the behavioral expression of learning and memory related performances [12]. However, the olfactory conditioning procedures engage usually many individuals at the same time e.g. in the T-maze paradigm ([13]), which makes it impossible to relate individual performances to their corresponding gene expression patterns. Olfactory conditioning protocols in *Drosophila* used by others also failed to decipher the heterogeneity in the expression of learned behavior among the individuals, rather the results were interpreted as the expression of probabilistic behavior at the individual level [14] [15]. Honeybees, on the other hand, can be selected for fast learning and then selectively bread via the thelytokous development of *Apis mellifera capensis* [16]; a highly effective procedure that should be applied more frequently. Tait and colleagues [17] tested individual honeybees in associative acquisition, non-associative sensitization, and short-term memory, and constructed a general heritable factor similar to factors derived for mammals [18] [19] [20] . The proboscis extension response (PER) training is a particularly effective experimental procedure that allows testing individual honeybees to a whole range of associative, non-associative and rule learning related paradigms in well controlled sequence of events [21] [22]. However, learning and retention scores are expressed in all these studies by data from groups of animals thus averaging across the individuals. Motivated by the study of Gallistel and coworkers [23] Pamir and coworkers [24] analyzed a large dataset on PER conditioning in the honeybee and found that once an animal started eliciting the conditioned response (CR), it kept showing the CR in the subsequent conditioning trials with a high probability. Importantly, the gradual acquisition in the groups of animals that results from the recruitment of individuals to the learner population, as predicted by Gallistel et al [23], was also found for the honyebees. Additionally, it has also been found that individual animals switched abruptly to a learned state during the acquisition phase after different number of acquisition trials and when they switched, they continued performing as learners Pamir [24]. Further, another study also reported that bees switching to learners early during the acquisition phase had higher retention scores later in the tests of memory retention but they did not discriminate better between the rewarded and unrewarded conditioned stimuli during differential conditioning [25].

These results have revealed some of the important aspects of learning behaviour of individuals which were previously unnoticed in the analyses of group-averaged learning behaviour albeit no clear picture has emerged on the different classes of bees which learn odours with different efficacy. We thus attempt to quantify this population heterogeneity by a sequential conditioning procedure that allows finer classification of individuals on the behavioral level based on their performances during two subsequent phases of differential olfactory conditioning and several retention tests, evaluating the strength of olfactory learning and sensitivity.

Multiple studies related different behavior in honeybees to their genetic backgrounds, such as the developmentally-regulated division of labor in the colony [26] [27], egg-laying and foraging behavior of the workers in queenless colonies [28], specialized tasks such as guarding and undertaking [29], foraging [30] [31], scouting and recruiting [32], as well as hygienic behavior against the *Varroa* mite [33]. It has also been shown that the manipulations of transcript levels in the honeybee brain influence memory performances [34] [35] [36]. Furthermore, a comparative transcriptomic study in the wasp, *Polistes metricus* revealed genetic associations of behavioral tasks and substantial overlap of genes involved in the division of labor in the honeybee and foraging in the wasp [37]. In the harvester ant, *Pogonomyrmex californicus,* large differences in gene expression, including the genes involved in neuronal functions and chemosensory perception, also have been reported for the queens showing different degrees of aggressive behavior [38]. We, therefore, selected the honeybees based on our previous work on their multiple forms of learning, their memory functions and the brain parts involved in these performances [1] [39], the mushroom bodies, which both in the bee brain as well as in *Drosophila* are known to be involved in higher order processing, learning and memory [40] [41] [42]. The aim of our study was to investigate the behavioral heterogeneity and whether it is associated with gene expression in the brain of the honeybee.

## Materials and methods

### Honeybee lines

A backcrossed genetic line of honeybees (*Apis mellifera carnica*), selected for improved hygienic behavior against the ectoparasitic *Varroa* mite, was used in this study (Länderinstitut für Bienenkunde (LIB), Hohen Neuendorf, Berlin [43]. This line was generated by artificially inseminating a queen from a non-hygienic (+/+) colony with the sperms from the hygienic drones (H). Heterozygous (H/+) queens were selectively raised from the F1 progeny and backcrossed with the sperms of their hygienic (H) paternal generation. These backcrossed heterozygous (H/+) queens were used to raise four colonies, (Colony 67, 73, 98 and 299) characterized by worker bees with homozygous (H/H) and heterozygous (H/+) genetic backgrounds for the hygienic trait. It is expected that these backcrossed colonies have more homogenous genetic backgrounds than natural colonies, which is advantageous for the behavioral and genetic investigations. Indeed, previous studies revealed that heterogeneity in olfactory learning performance of honeybees has genetic basis [44] [45] [46].

The experiment took place between July and October 2010. During the course of our experiment, colony 67 became aggressive and was quarantined. From the time point of its confinement, we have excluded bees from this colony from the data analysis (a total of 32 bees were excluded).

### Procedure for sequential olfactory PER conditioning

Forager honeybees were caught at the entrance of the colonies on the day before sequential conditioning. Bees were harnessed in small plastic tubes as described elsewhere [21] [47] [48], fed until satiation with 30% (W/V) sucrose solution and kept overnight inside a humid Styrofoam box in the dark. The next morning (day-2), they were placed in front of the experimental arena for 30–45 min for adaptation. The sequential conditioning protocol consisted of two identical phases, each composed of one round of differential conditioning followed by two sets of retention tests (Fig 1). During the differential conditioning, bees were trained to learn and discriminate an odor (conditioned stimulus, CS+) paired forwardly with the presentation of a sucrose reward (a 30% sucrose solution was used as unconditioned stimulus, US) from a non-reinforced odor (CS–) in the course of 12 consecutive conditioning trials (6 CS+ and 6 CS–). The two pure odor stimuli were presented alternately starting with the CS+ with an inter-trial interval (time gap between consecutive CS+ and CS– trials) of 8 min. During the CS+ trials, the US was applied 3 s after the odor onset by first touching the antenna with a toothpick soaked with the sucrose solution to elicit proboscis extension, followed by feeding through the proboscis for a total of 4 s. During the CS– trials and the retention tests, the odors were presented alone for 5 s. During conditioning, a PER elicited during the first 3 s of odor stimulation before the onset of the US was considered as a conditioned response (CR). A 20 min break was given after the differential conditioning, which followed two sets of retention tests. In each test, bees were presented with increasing concentrations of the CS+ and CS– odors on filter paper (10^-3^ and 10^-2^ dilutions in paraffin oil and the pure odors) starting with the CS+. During each set of retention test, the animals were also stimulated once with paraffin oil and once with the filter paper alone. There was no time gap between the two consecutive sets of retention tests. The second phase began after a pause of 30 min and a different odor pair was used for the second differential conditioning (Fig 1). The ability of the bee to elicit a CR was evaluated by showing that the stimulation of the antennae with sugar water elicited PER. Only bees that elicited PER were considered for data analysis. Bees were sacrificed immediately after this fitness-test, by placing them at first –20°C for 10 min followed by preservation at –80°C until the gene expression study. The whole procedure ran for a period of 6 h and 30 min.

**Fig 1:**
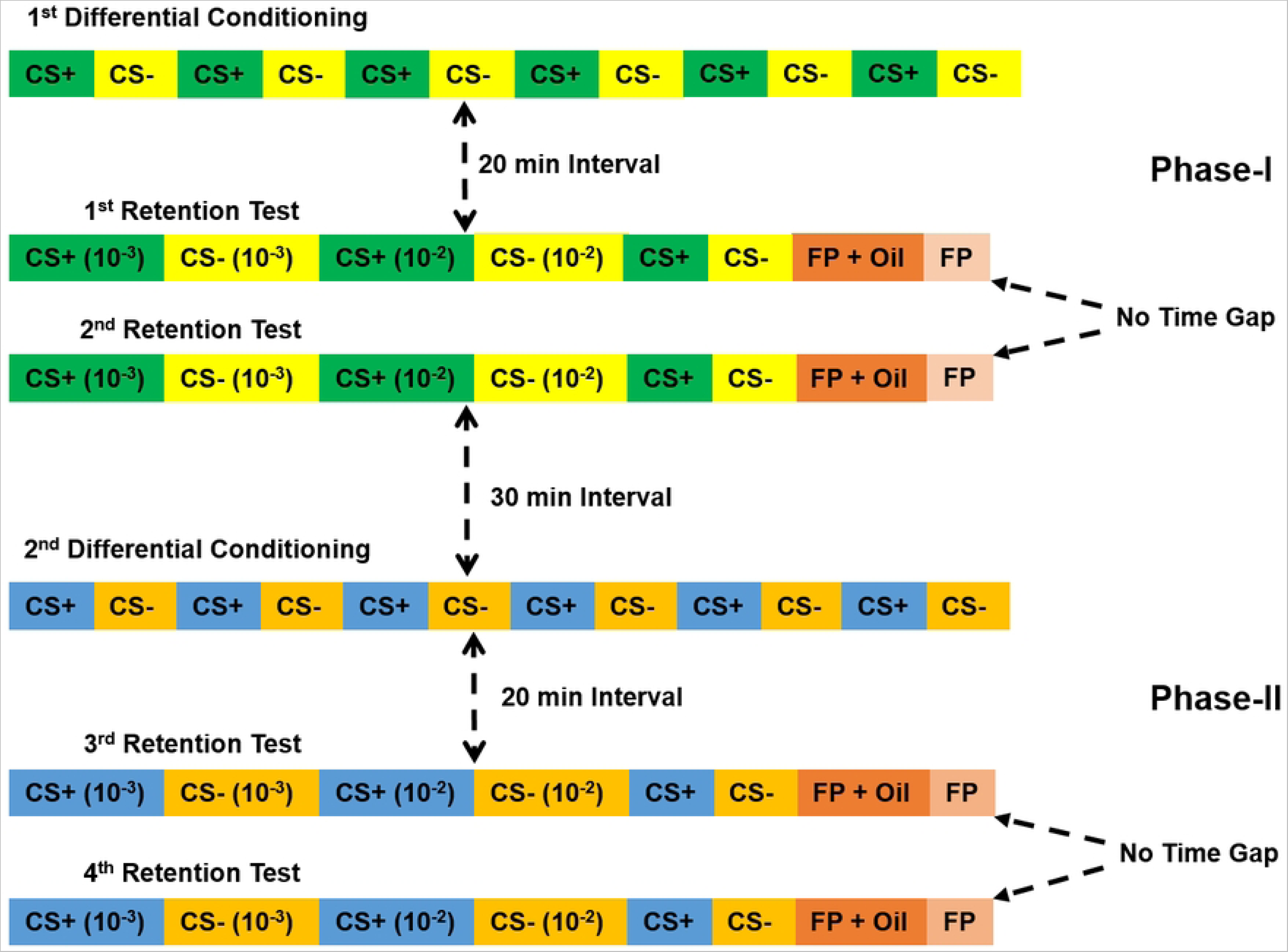
The protocol of sequential conditioning includes two phases each composed of one differential olfactory conditioning and two sets of retention tests. Different odors were used (color-coded) for the two phases and in the tests, dilutions of the odors were also used along with the pure concentrations, used for conditioning. Responses to the filter paper and paraffin oil were also recorded. The time intervals between the conditioning and tests and between the two phases are mentioned in the figure.

Natural odors were used as CSs. During the first phase, we used the volatile odors specific for the healthy brood (β-ocimene or OM, purity >90%, Sigma Aldrich, Germany) and larvae infected by chalkbrood pathogen, *Ascosphaera apis* (phenethyl acetate or PEA, purity 99%, Sigma-Aldrich, Germany) (Swanson et al., 2009). It was reported that hygienic or *Varroa*-tolerant bees discriminate well between these odors [49] [50] [51]. Two volatile fatty acids, oleic acid (OA, purity >99%, TCI-Europe NV, Belgium) and linolenic acid (LA, purity >70%, TCI-Europe NV, Belgium), reported to be involved in the process of kin-discrimination in honeybee [52] were used as odors for the second phase of sequential conditioning. They have similar chemical structures and thus were expected to be harder to differentiate. During the two phases of conditioning, odor contingencies were reversed to understand whether each odor can be learned well either as a CS+ or as a CS-by the bees. Odor stimuli were delivered with a 20 ml plastic syringe containing a filter paper soaked with 10 µl of the respective odor. The odor stimuli were delivered manually for a period of 5 s.

### Analysis of behavioral data – Quantification of an individualized performance score

A total of 171 honeybees were included in the data analysis. We scored the following seven behavioral parameters to evaluate the olfactory learning performance of the individual honeybees in our assay: Feature-1 and -2: Rate and reliability of CS+ odor learning during the acquisition trials of the first (Acq-1) and the second (Acq-2) differential conditioning: The CR was normalized to the CS+ for individual bees from the 2^nd^ to the 6^th^ trial during the differential conditioning. The score ranges from 0 to +1. Feature-3 and -4: Odor discrimination during the acquisition trials of the first (DisCond-1) and the second (DisCond-2) differential conditioning trial. Normalized CS+ responses that were not followed by a response to the CS– from the 2nd to the 6^th^ acquisition trial during the differential conditioning. The scores ranged from 0 to +1. For feature-1 to -4, we only scored the responses of the bees starting from the second conditioning trial of the CS+ and CS– because a response was considered to be a conditioned response only after the first conditioning trial. Feature-5 and -6: Odor discrimination during the first and second (DisT-1,-2) sets of retention tests and the third and fourth (DisT-3,-4) sets of retention tests: Normalized responses to the CS+ stimuli (2 dilutions and pure odors) during the retention test that were not followed by a response to the CS– stimuli (2 dilutions and pure odors). Thus, the scores ranged from 0 to +1. An individualized cumulative or overall performance score (P-score) was calculated for each bee by adding the scores for the six behavioral parameters. The score ranged from 0 to +6.

### Statistical analysis of behavioral data

Conditioned responses of the bees were analyzed by repeated measures ANOVA (RM-ANOVA) followed by the Tukey HSD posthoc test [53] [54] [55] [56]. Wilcoxon Matched Pairs test was applied to compare the scores of the behavioral features for the same set of bees between the two phases. The total number of PER to the CS stimuli during the conditioning and retention tests was compared between the two phases using the G-test. Lilliefors test was performed to understand the nature of distribution of P-scores in the data. Mann-Whitney *U* test was used to compare the scores of behavioral features between the high and low performing bees. Pearson’s linear correlation coefficients were calculated between the scores of the six parameters to understand the nature of associations between them. Linear regression analysis was performed separately for each of the six parameters to find out how much of the variability in P-score is explained by each of them. The programs used for statistics and figure-making were Statistica version 5.0 (Dell Software, USA), SPSS version 23 (IBM), and MatLab (MathWorks, USA). The analyses of significant gene overlap between the present study and [57] were performed with a χ^2^ test (www.socscistatistics.com/tests/chisquare/default2.aspx). The test was applied by considering that the honeybee genome comprises 13440 genes. The differences were considered statistically significant when *p* < 0.05.

### Tissue preparation and total RNA extraction

The head capsule of the bee was separated from the rest of the body and a window was cut in the cuticle. The manipulation was performed on a metal slide cooled on dry ice. The head was partially lyophilized in a vacuum chamber at –20°C for 2 hours and then fixed with Tissue-Tek on a metal slide cooled on dry ice. The brain was exposed by scraping off the hypopharyngeal glands covering the brain and the dorsal part of the central brain containing the mushroom body was recovered and immediately homogenized in 400 µl Trizol (Life Technologies, Schwerte, Germany) in a Teflon glass homogenizer. After phase separation, the aqueous phase containing the total RNA was recovered, mixed in equal volume with 100% ethanol, loaded on a column (RNeasy MinElute Cleanup, Qiagen, Hilden, Germany), treated with DNAse (Qiagen, Hilden, Germany), washed and eluted in water.

### Microarray analysis

The methodology described by Gempe and colleagues was applied [33]. The honeybee whole-genome oligonucleotide microarray (Design: UIUC Honey bee oligo 13 K v1, Accession: A-MEXP-755), containing 28,800 oligos that represent 13,440 genes was derived from annotations of the entire honeybee genomic sequence [58]. Ten microliters of each of the total RNA sample was amplified prior to labelling following the manufacturer’s protocol (MessageAmp II aRNA Amplification kit, Ambion). We hybridized 3 μg of each labelled RNA sample to a single microarray slide. The slides were scanned (SureScan Microarry scanner, Agilent, Waldbronn, Germany) and raw hybridization signals were extracted (GenePix Pro 6.0 software, Agilent Technologies). Transcription-level data were processed and analyzed using the LIMMA 2.16 software package (https://www.bioconductor.org/packages/3.3/bioc/html/limma.html). The quality of hybridization was evaluated using the raw expression data from control probes spotted on each slide. Transcription-level data were corrected for background signal (“normexp” function) [59] and intensity-dependent bias was detected (“normalize within arrays” function with the default print-tip lowess normalization method) [60]. Finally, the log-transformed expression ratios were calculated. Data from duplicate spots were averaged using the “avedups” function. We used a design matrix that incorporated the P-score, the colony conditions and linear models using the Bayesian fitting option. All microarray data are MIAME-compliant, and the raw data have been deposited in a MIAME-compliant database (ArrayExpress, EMBL-EBI – https://www.ebi.ac.uk/arrayexpress/experiments/E-MTAB-7909/). Differences in gene transcription that resulted from behavior or from the type of backcross were specified as separate contrasts using linear models. P-values were adjusted for multiple testing with a 5% false discovery rate. Functional annotation of gene sets that fell into similar categories of GO terms for molecular processes and biological functions were identified using DAVID (http://david.ncifcrf.gov/) [61], [62], and KEGG [https://www.genome.jp/kegg/], which includes an enrichment analysis of GO terms. We used the gene annotations from the UIUC Honey Bee oligo 13 Kv1 annotation file.

### RT-PCR

In addition, confirmatory expression analyses were also performed for selected transcripts using RT-qPCRs according to the previously published protocols. [63] [64]. Primers are listed in Supplementary Table 1 (S1) and *Actin* was chosen as reference gene (TIB Molbiol, Berlin, Germany). Fold change was calculated using the 2^-ΔΔCt^ method. Data points deviating by more than 2-standard deviations to mean were excluded. Statistical differences were evaluated using Student’s t test for independent measures.

## Results and discussion

We have trained two independent groups of bees in the sequential conditioning paradigm along two phases with each phase comprising one differential conditioning and two sets of retention tests, involving two pairs of odor; PEA and OM were used for the first phase, LA, and OA for the second phase. To balance odor contingencies, Group-1 (N = 85 bees) bees were trained to discriminate between PEA as the rewarded odor (CS+) and OM as the unrewarded odor (CS-) in the first phase of sequential conditioning while in the second phase, LA was rewarded (CS+) and OA was unrewarded (CS-) (Table 1). While, Group-2 (N = 86 bees) was conditioned with OM as CS+ and PEA as CS-, followed by OA as CS+ and LA as CS-. We first compared the performances of the two groups during the two differential conditioning and found no significant difference between Group-1 and Group-2 within each phase, as revealed by 2 × 2 × 6 (group × stimulus CS+/CS- × trial) ANOVA for repeated measures (Table 2).

**Table 1:**
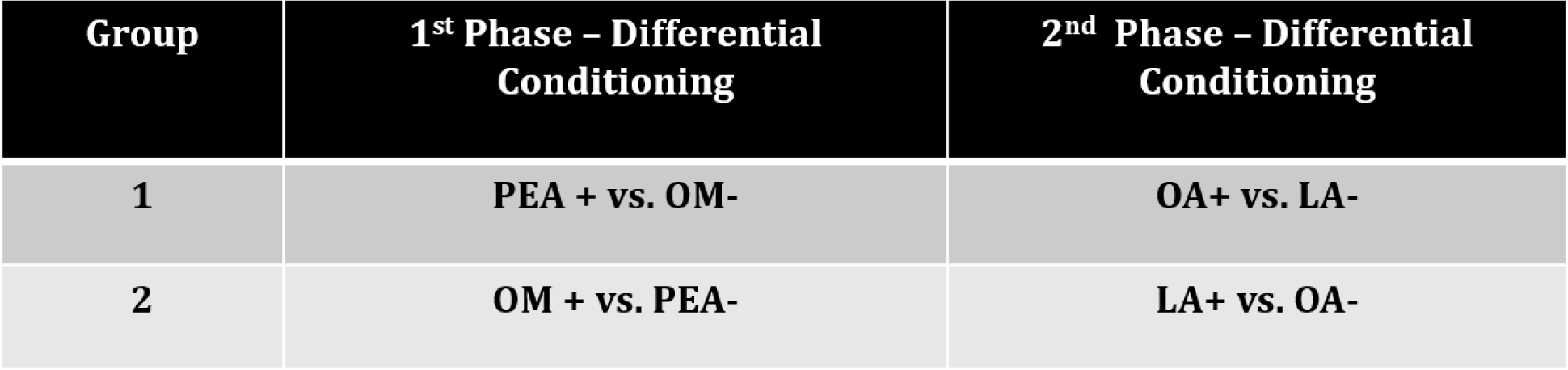
Two separate groups of bees were trained along two differential conditioning in the sequential conditioning paradigm. Phenethyl acetate (PEA) and β-ocimene (OM) were used as CS odors in the first conditioning and oleic acid (OA) and linolenic acid LA) were used in the second conditioning. The contingencies (+: reinforced, -: unreinforced) of the odors were reversed for the two groups of bees for each of the two differential conditioning to keep the balance in odor contingency.

**Table 2:** Performances of Group-1 and -2 were compared using ANOVA test for repeated measures within the individual differential conditioning. Fisher statistic values given in the table for the main effect (Group) and the three other interaction effects are all statistically non-significant (*p* > 0.01). Thus, performance of both groups were pooled.

| Main and Interaction effects | 1<sup>st</sup> Phase – Differential Conditioning | 2<sup>nd</sup> Phase – Differential Conditioning |
| --- | --- | --- |
| Group | $F_{1,338} = 1.67$ | $F_{1,338} = 0.0003$ |
| Group × Stimulus | $F_{1,338} = 2.28$ | $F_{1,338} = 0.48$ |
| Group × Trial | $F_{5,1690} = 1.49$ | $F_{5,1690} = 2.14$ |
| Group × Stimulus × Trial | $F_{5,1690} = 1.33$ | $F_{5,1690} = 0.3$ |

We then compared the performance of the two groups in the four sets of retention tests and found a significant interaction effect between trial and group (F_2,676_ = 23.28, *p* < 0.01) only for the first of the two sets of retention tests of the first phase of sequential conditioning. This significant interaction effect was due to an overall higher response of Group-2 than Group-1 only during the first odor trial of the first retention test because for rest of the two test trials, the two groups showed similar responses to the different concentrations of the CS odors. We found no significant interaction effects for the set of first retention test; group and stimulus (F_1,338_ = 2.61, *p* > 0.05), group, stimulus and trial (F_2,676_ = 2.57, *p* > 0.05). No significant main effect and interaction effect was also found for the second set of retention test (data not shown). Similar to the first set of retention test, significant interaction effects were also found between trial and group for both the third (F_2,676_ = 44.61, *p* < 0.01) and fourth set of retention tests (F_2,676_ = 12.87, *p* < 0.01), however no significant interaction effects (between group and stimulus, and between group, stimulus and trial) were found for these two sets of retention tests; third set of test: group × stimulus (F_1,338_ = 2.21, *p* > 0.05), group × stimulus × trial (F_2,676_ = 0.85, *p* > 0.05), fourth set of test: group × stimulus (F_1,338_ = 0.48, *p* > 0.05), group × stimulus × trial (F_2,676_ = 0.09, *p* > 0.05). Since no significant differences were found in the overall responses of the bees of the two groups to the CS odors during the two differential conditioning and four sets of retention tests (non-significant interactions between group, stimulus and trial), we pooled the data of the two groups for all further analyses.

### Learning and memory performance of the honeybees in the sequential conditioning

In the first differential conditioning, bees learned the discrimination task (CS+ vs. CS-) successfully. A 2 × 6 (group × stimulus) ANOVA for repeated measures produced significant stimulus and trial effects as well as a significant interaction effect (F_5,1700_ = 99.32, *p* > 0.01) (Table 3). Similarly, significant stimulus, trial and stimulus × trial effects are also found for the second differential conditioning (Table 3). Thus, the results of the repeated measures ANOVA test demonstrate that bees learned the discrimination task well between the CS+ and the CS– odors during the two differential conditioning (Fig 2A, 2F). The response levels to the CS+ increased across the conditioning trials as the responses to the CS– decreased. The differences in response levels to the CS+ and CS– increased across the successive trials of the four sets of retention tests, which correlates with the increasing odor concentrations of the CS odors from the first to the third trial of the retention tests (Fig 2B, 2D, 2G, and 2I, Supplementary Table 1). During the fourth set of retention test, bees could not discriminate between the CS+ and CS- at their lowest concentration (10^-3^) (Table S2, Fig 2I). The response levels diminished for the CS– during the retention tests to reach the response levels of the filter paper and paraffin oil tests (Fig 2C, 2E, 2H, and 2J). No significant differences were found between the responses to the filter paper alone and filter paper soaked with paraffin oil for the two phases of sequential learning (data not shown).

**Fig 2:**
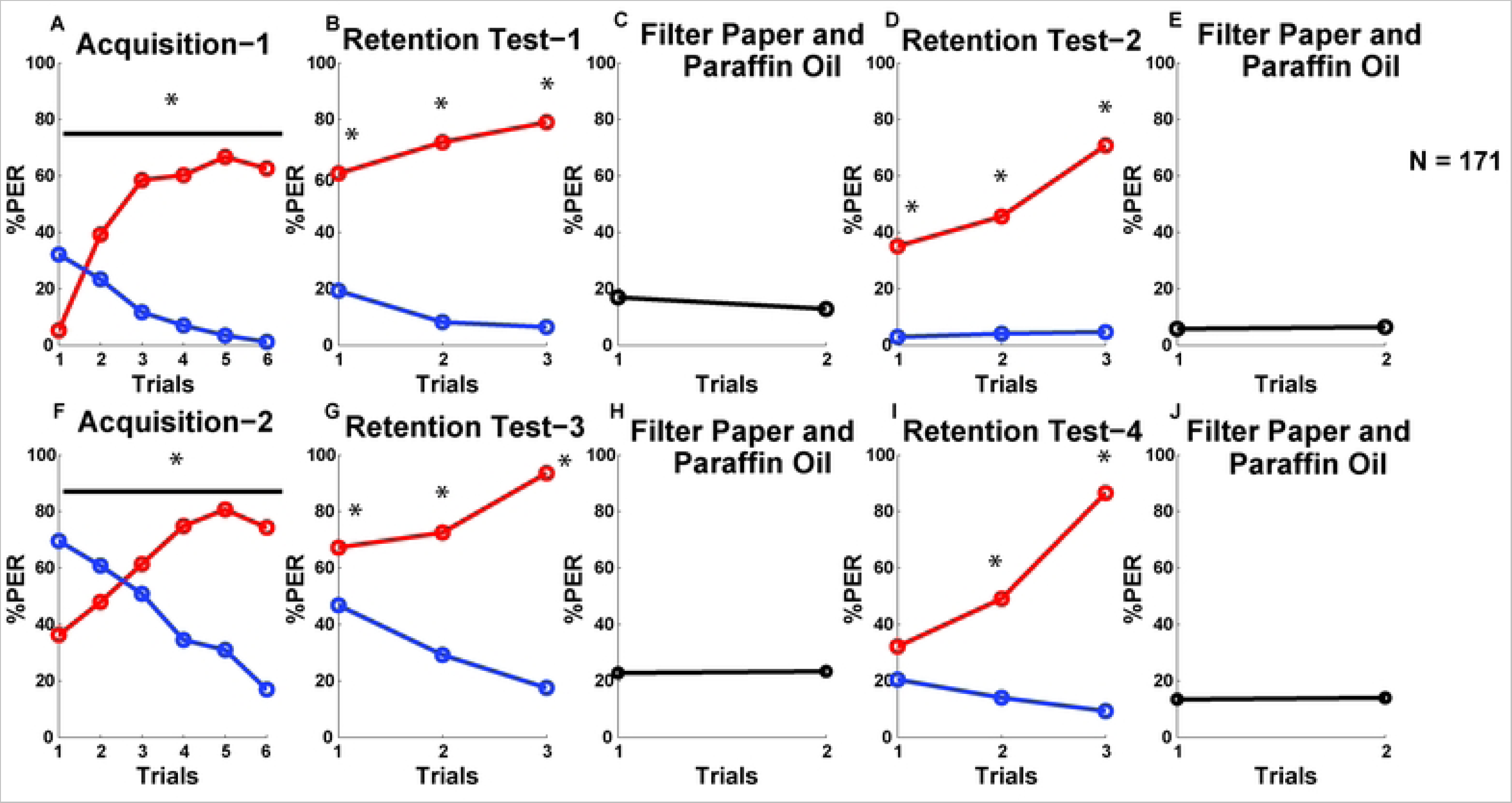
Line graphs, showing the percent PER to the CS+ (red line) and CS- (blue line) stimuli for the pooled population (N = 171) during the two differential conditioning (A and F) and four sets of retention tests (B, D, G, and I) of the sequential conditioning. Responses to filter paper soaked with paraffin oil (first data point) and filter paper only (second data point) are shown in C, E, H and J. Asterisks are denoting statistically significant differences (*p* < 0.01) between the PERs to the CS+ and CS– odors. See Table 3 and Table S2 for statistics.

**Table 3:** Fisher statistic values (pooled data) from the repeated measures ANOVA tests are given for the two differential conditioning (pooled data). All *p*-values correspond to the stimulus, trial and stimulus × trial effects are significant (*p* < 0.01).

| Phase of Sequential Learning | Stimulus Effect | Trial Effect | Stimulus × Trial Effect |
| --- | --- | --- | --- |
| 1<sup>st</sup> | $F_{1,340} = 164.65, p < 0.01$ | $F_{5,1700} = 12.39, p < 0.01$ | $F_{5,1700} = 99.32, p < 0.01$ |
| 2<sup>nd</sup> | $F_{1,340} = 30.86, p < 0.01$ | $F_{5,1700} = 3.76, p < 0.01$ | $F_{5,1700} = 81.96, p < 0.01$ |

### Differential responses during the two phases of sequential conditioning

We found honeybees that underwent sequential conditioning learned the odor stimuli but they showed differential responses in the two phases. To investigate the differences in PER responses between the first and second phase, we compared the odor discrimination scores quantified for the acquisition trials of the first (DisCond-1) and the second (DisCond-2) differential conditioning. We found that bees showed significantly higher odor discrimination in the first than the second differential conditioning (Wilcoxon Matched Pairs test: Z = 4.11, T = 2715, *p* < 0.01) (Fig 3A). This is because bees showed significantly higher total number of responses to the CS- odors during the second than the first differential conditioning (first Phase: 491 PERs to the CS+ and 80 PERs to the CS-, second Phase: 580 PERs to the CS+ and 332 PERs to the CS-) (G-test: G = 214.8, df = 1, *p* < 0.01). When we compared odor discrimination scores between the retention tests of the first phase and the second phase of sequential conditioning, we also found a significantly higher odor discrimination during the first and second (DisT-1,-2) sets of retention tests than the third and fourth (DisT-3,-4) tests (Wilcoxon Matched Pairs test: Z = 2.42, T = 3543, *p* < 0.01) (Fig 3B), which confirms that bees overall discriminated better between the dilutions of the CS+ and CS– odors in the first and second sets of the retention test than in the third and fourth sets of test. Likewise differential conditioning, bees also showed significantly higher total number of PERs to the CS- odors during the 3^rd^ and 4^th^ sets of retention test than the first and second sets of test (first + second retention tests: 621 PERs to the CS+ and 78 PERs to the CS-, third + fourth retention tests: 686 PERs to the CS+ and 235 PERs to the CS-) (G = 96.63, df = 1, *p* < 0.01). These results altogether show that bees mastered the odor discrimination tasks more efficiently in the first half of the sequential conditioning procedure and generalized more during the second half. Importantly, the odors used in the second half were also relatively harder to discriminate, especially when diluted (see Fig 1 and Table S2).

**Fig 3:**
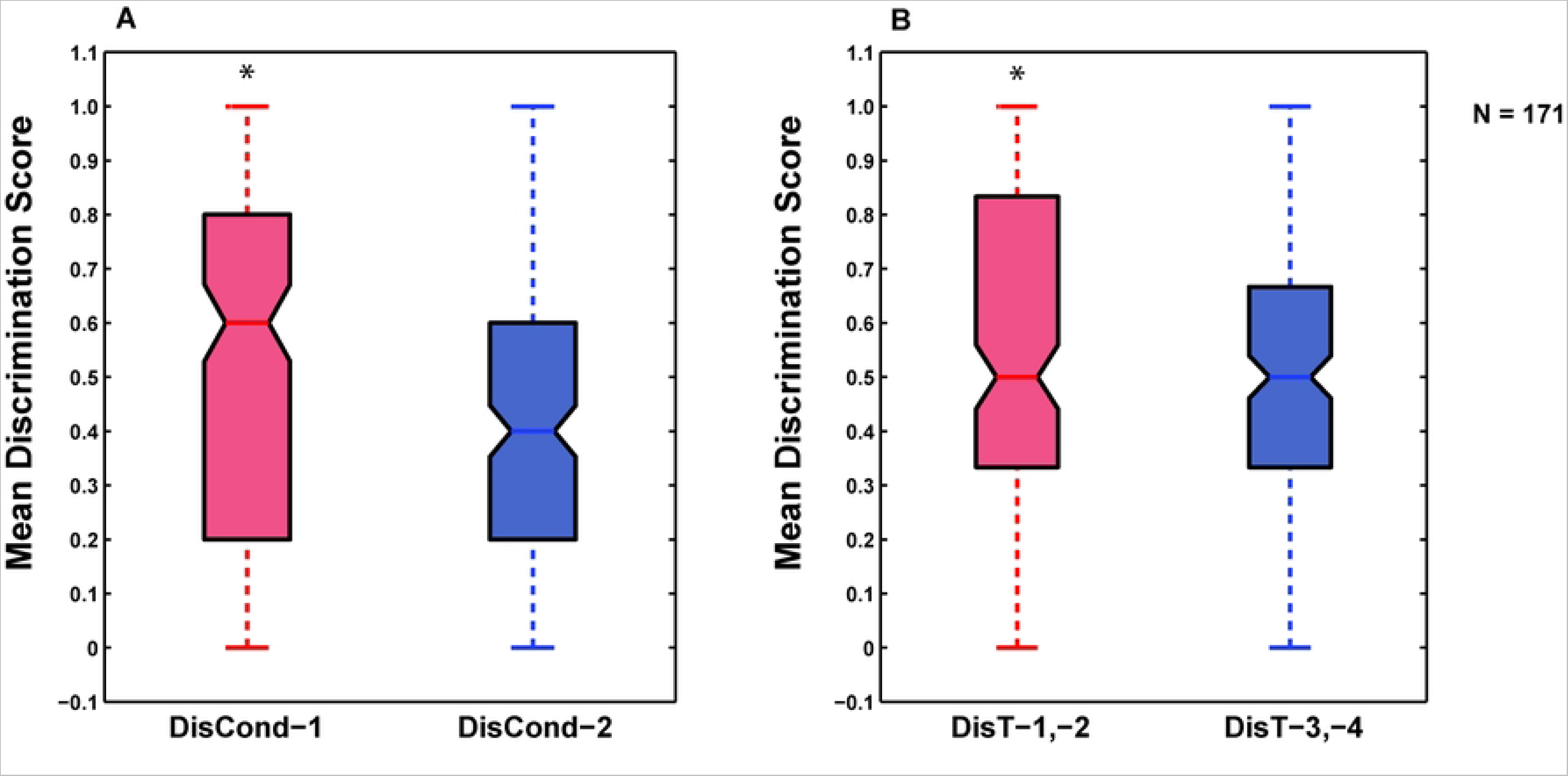
Comparison of the discrimination scores between the CS+ and the CS- during the conditioning phases (A) and the retention tests (B). The box and whisker plots show the distributions of odor discrimination during the first (DisCond-1, red) and the second (DisCond- 2, blue) conditioning phases (A) and during the first and second (DisT-1,-2; red), and the third and fourth (DisT-3,-4; blue) retention tests (B). Asterisks denote statistically significant differences (*p* < 0.01). The number of animals is indicated (N).

### Classification of the honeybees into various performance categories based on their individual P-scores

Overall performance levels of the individual honeybees in the sequential conditioning procedure are represented by a cumulative performance score (P-score), which is the summation of scores of the six quantified parameters (Fig 4). The distribution of P-scores is found to differ significantly from a Gaussian distribution (Lilliefors test: *p* < 0.01) (Fig 4). The P-score of bees varies between 0 and 5.46 with a mean score of 3.12 +/- 1.34 SD. Only 27 bees (∼15.8%) scored ≤ 1.78, (< 1-SD of the mean), and 28 bees (∼16.4%) scored ≥ 4.46(> 1-SD of the mean) whereas the majority of bees (116 bees, ∼67.8%) had scores between 1.78 and 4.46, corresponding to +/- 1-SD of the mean score. We compared the performance levels of the bees with low (P-score<2.6) and high P-scores (P-score>=2.6), which we respectively categorized as low and high performers. The acquisition and retention test performances are substantially better in the high performer bees than the low performers all-round the two phases of sequential conditioning (Fig 5, Table 4). The low performer bees not only showed an overall reduced number of responses to the CS odors throughout the procedure but also failed to discriminate between the pure odors during the first differential conditioning as well as between their dilutions during the retention tests-2 and -4 (Fig 5; see Table S3 for the statistical analysis of within-group performance). We applied Mann-Whitney *U* test and compared the scores of all of the six parameters viz., Acq-1, Acq-1, DisCond-1, DisCond-2, DisT-1,-2, and DisT-3,-4 of these two categories of bees. The results show that high performer bees scored significantly higher than the low performers for all of the six parameters (Table 4). We also separately analyzed the responses of these two groups of bees to the novel stimuli, filter paper (FP) and paraffin oil (Oil), which were used in the retention tests. We found that high performers also showed significantly higher total number (altogether in the two phases of the procedure) of responses to these stimuli than the low performer bees (G =24.48, df = 1, *p* < 0.001) (responses to FP and Oil are shown in Fig 5, Table 4).

**Fig 4:**
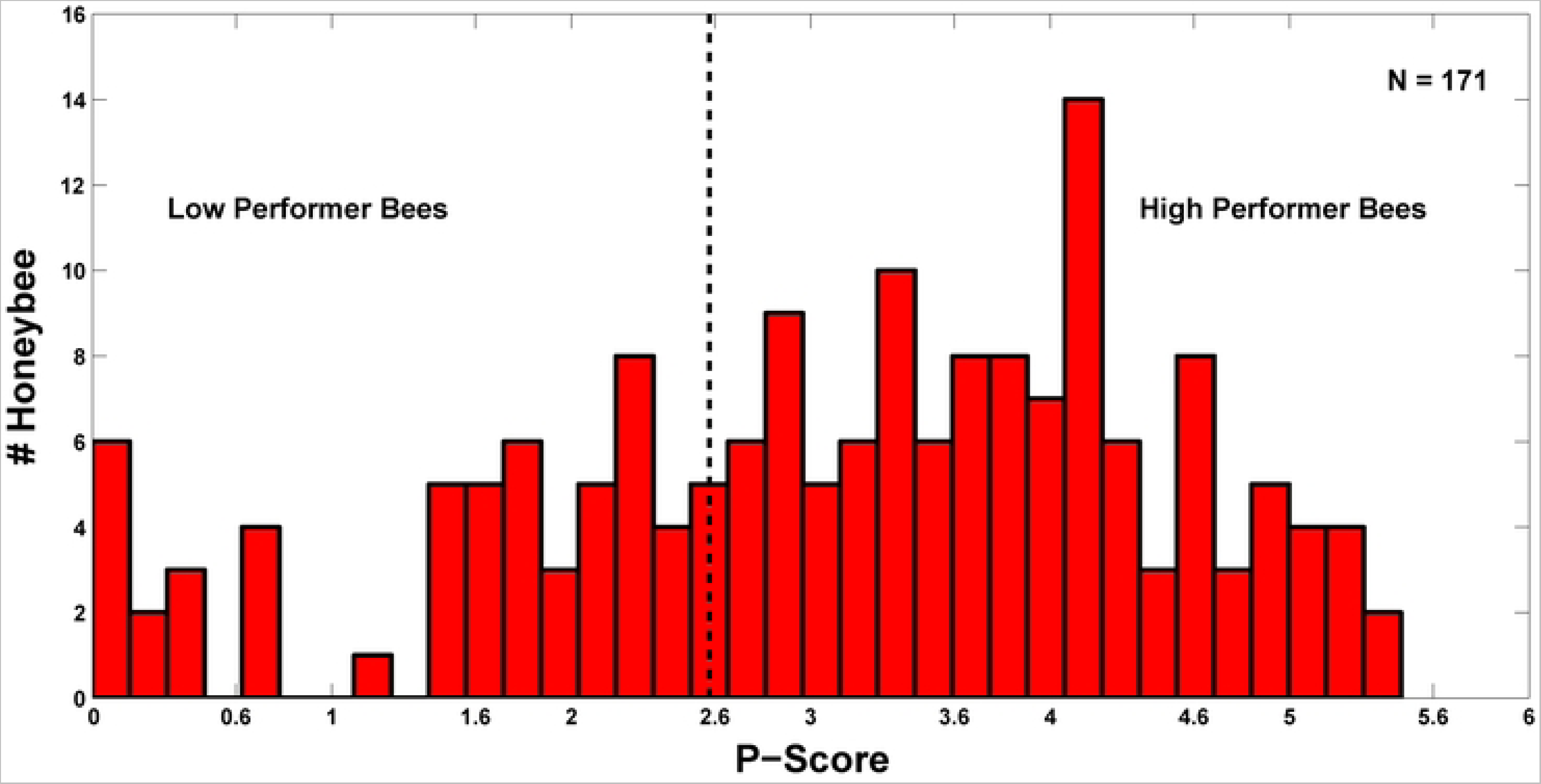
Histogram of P-scores for the pooled data of bees. The dotted black line represents the threshold in P-score (2.6), used for the selection of low performer (P-score<2.6) and high performer bees (P-score=>2.6). N represents total number of animals.

**Fig 5:**
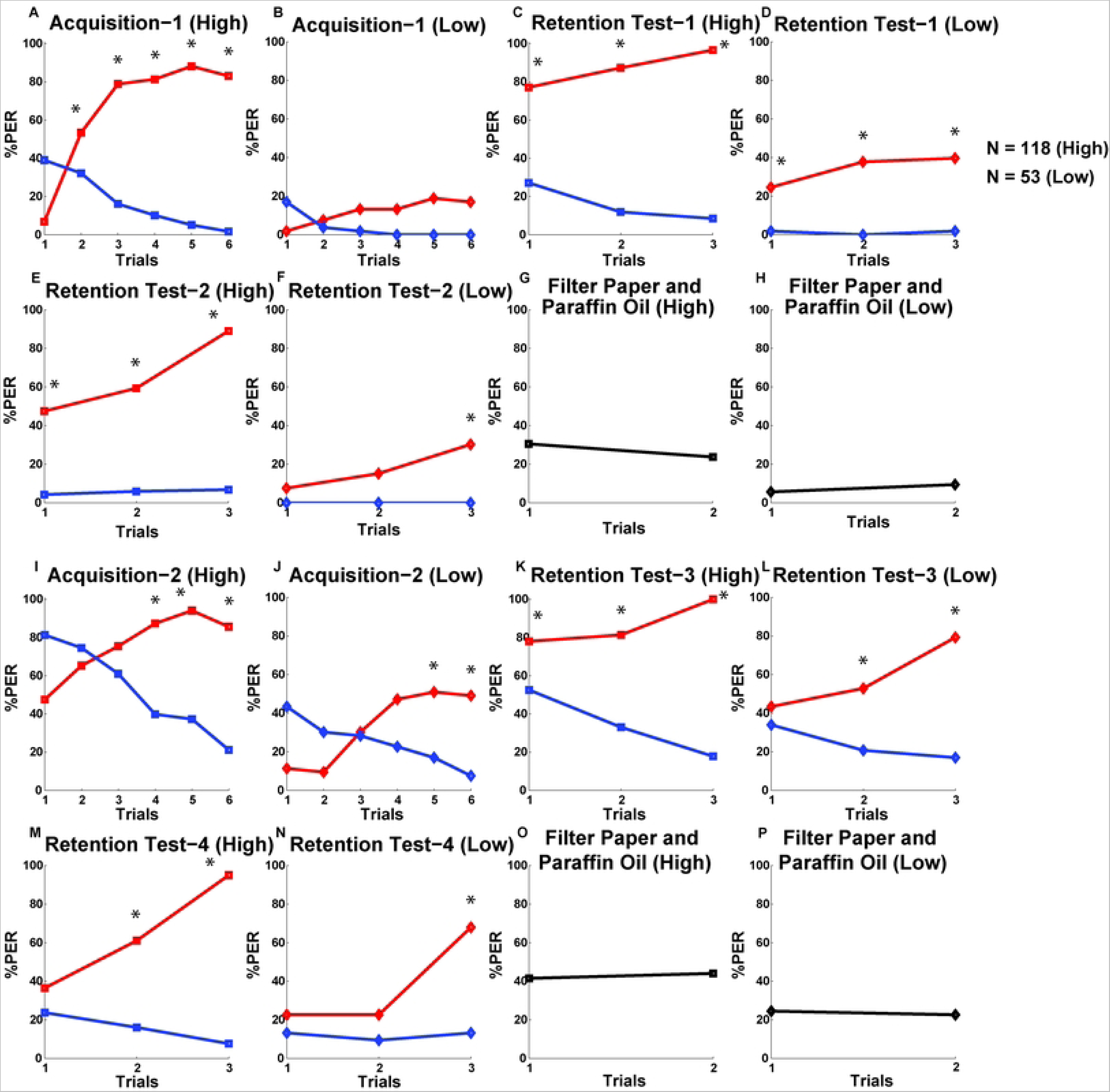
Line graphs, showing the percent PER to the CS+ (red line) and CS- (blue line) stimuli for the high performer (N = 118) and low performer (N = 53) bees during the two phases of sequential conditioning and retention procedures. The first set of eight subfigures (A to H) represent the PERs to the odors during the first phase. Line graphs of the high and low performers for the same odor trials are given in a pairwise manner, such as Acquisition-1 (High and Low), Retention Test-1 (High and Low), Retention Test-2 (High and Low), Filter Paper and Paraffin Oil (High and Low) to visualize the differences in performance between these two groups of bees. The arrangement of the line graphs are the same for the two groups of bees for the second phase of sequential learning (Subfigure I to P). Total number of responses to the filter paper soaked with paraffin oil (first data point) and filter paper only (second data point) for a particular phase (first phase: summation of responses the first and second = retention tests, second phase: summation of responses for the third and fourth retention tests) are given for these two groups of bees (Subfigure G, H, O, and P).Repeated measures ANOVA followed by Tukey’s HSD posthoc test were performed to analyze the performances of the two groups of bees during the conditioning and retention stages (see Table S2). Significant differences (after Bonferroni correction) in responses are represented by asterisks.

**Table 4:** Results of the Mann-Whitney *U* test are showing that high performer bees (N = 118) scored significantly higher than the low performers (N = 53) in all of the six quantified parameters related to odor learning and discrimination during the two differential conditioning and four sets of retention tests of the sequential learning paradigm. The table shows the mean scores of each of the six parameters for the two selected groups of honeybees.

| Parameter | Comparisons Between High and Low Performer Bees |
| --- | --- |
| **Acq-1** | (Mean Acq-1<sub>high</sub> = 0.76, Mean Acq-1<sub>low</sub> = 0.13)<br>Z-value = 9.94, $p < 0.001$ |
| **Acq-2** | (Mean Acq-2<sub>high</sub> = 0.81, Mean Acq-2<sub>low</sub> = 0.37)<br>Z-value = 7.67, $p < 0.001$ |
| **DisCond-1** | (Mean DisCond-1<sub>high</sub> = 0.66, Mean DisCond-1<sub>low</sub> = 0.13)<br>Z-value = 9.49, $p < 0.001$ |
| **DisCond-2** | (Mean DisCond-2<sub>high</sub> = 0.42, Mean DisCond-2<sub>low</sub> = 0.25)<br>Z-value = 3.85, $p < 0.001$ |
| **DisT-1,-2** | (Mean DisT-1,-2<sub>high</sub> = 0.65, Mean DisT-1,-2<sub>low</sub> = 0.25)<br>Z-value = 7.54, $p < 0.001$ |
| **DisT-3,-4** | (Mean DisT-3,-4<sub>high</sub> = 0.52, Mean DisT-3,-4<sub>low</sub> = 0.33)<br>Z-value = 4.60, $p < 0.001$ |

These results demonstrate that high performer bees performed consistently better than the low performers throughout the two phases of sequential learning and retention procedures. The high performers showed early and consistent responses to the CS+ odors and discriminated well between the CS+ and CS- odors during the two differential conditioning, as well as successfully discriminated between the different concentrations of the CS+ and CS- odors during the four sets of retention tests. In addition, high performer bees were also probably more sensitive and/or aroused than the low group owing to their higher responsiveness to the novel odors of filter paper and paraffin oil.

### Predictor analysis of the P-score

Next, we investigated how well the scores of the six quantified parameters, related to olfactory learning and memory, predict the P-score. We first looked at the linear correlation between these six parameters and found a high correlation (r = 0.91) between the speed of acquisition of the CS+ (Acq-1) and odor discriminability (DisCond-1) during the first phase of differential conditioning (Table S4). The speed of acquisition of the CS+ and odor discriminability during the first phase also show high correlation coefficient values with the P-score (respectively r = 0.85 and r = 0.84) (Table S4). The parameter, Acq-2 shows the next highest correlation with the P-score (r = 0.76) (Table S4).

To understand the capacity of each of the six parameters to predict the P-score, we performed linear regression analyses to find out how much of the variability in P-score is explained by each of the parameters separately. Due to significant correlations between some of these six parameters (Table S4), regression analysis was done separately for each parameter that were treated as independent variable, to avoid the influence of one over the others. Regression analysis shows that Acq-1 alone explains the maximum amount of variance (73.3%) in the P-score with each unit-score increase of Acq-1 leading to an increase of 3.18 unit-score of the P-score (B = 3.18) (Table 5). Like, Acq-1, DisCond-1 also explains a substantial amount of variance (71.3%) in the P-score (Table 5). However; the quantification of DisCond-1 includes the number of CS+ responses during the first differential conditioning, which is the measure for Acq-1. Thus, DisCond-1 depends on Acq-1 in predicting the P-scores. The other four parameters also contribute to the variance of the P-score: Acq-2 (57,9 %), DisT-1,-2 (55,9 %), DisT-3,-4 (37,2 %) and DisCond-2 (26,3 %) (Table 5). These results altogether illustrate that the rate of learning of the CS+ odors during the first phase of differential conditioning or Acq-1 has the highest predictive capacity for the P-score.

**Table 5:** Results of the regression analysis shows that Acq-1 explains the highest, 73.3% variance in the P-score. The other five parameters explain lesser amounts of variance. All model statistics are found significant (*p* < 0.01).

| Parameters | R<sup>2</sup>-(coefficient of determination) value | Model statistics (ANOVA) | Unstandardized regression coefficient (B) |
| --- | --- | --- | --- |
| Acq-1 | 0.733 | $F_{(1,169)} = 463.82, p < 0.001$ | 3.18 |
| Acq-2 | 0.579 | $F_{(1,169)} = 232.5, p < 0.001$ | 3.18 |
| DisCond-1 | 0.713 | $F_{(1,169)} = 420.45, p < 0.001$ | 3.45 |
| DisCond-2 | 0.263 | $F_{(1,169)} = 60.33, p < 0.001$ | 2.59 |
| DisT-1,-2 | 0.559 | $F_{(1,169)} = 214.49, p < 0.001$ | 3.29 |
| DisT-3,-4 | 0.372 | $F_{(1,169)} = 100.25, p < 0.001$ | 3.37 |

### Microarray analysis of high and low performer bees

Twenty-four bees were selected for the microarray analysis based on their performance during the behavioral tests, the colony and the date of the experiment (Table 6, Fig 6). Bees from the four colonies 67, 98, 73 and 29 were used for this analysis. The total RNAs extracted from the mushroom bodies were processed to generate the microarray probes. Two sets of probes were generated for each bee, one labelled with Cy5 and the other with Cy3. In this manner, each animal was analyzed twice by swapping colors on each array, to reduce artefacts due to the dye. The last bee of the list was analyzed with the first one, thus the analysis scheme was designed to build a loop. High performer bees had P-scores ranging between 2.6 and 5.4 and low performers between 0 and 2.5 (Fig 6). Eight bees from the colonies 98 and 73 and four bees from the colonies 67 and 299 were analyzed (Table 6). To reduce the colony effects on the identification of genes, high performer bees with P-scores ranging from 3.9 to 5.3 from the four colonies were also compared on 12 arrays (Table S4). In addition, each colony was also analyzed with three bees collected from July to October and dye swap loop design was applied.

**Fig 6:**
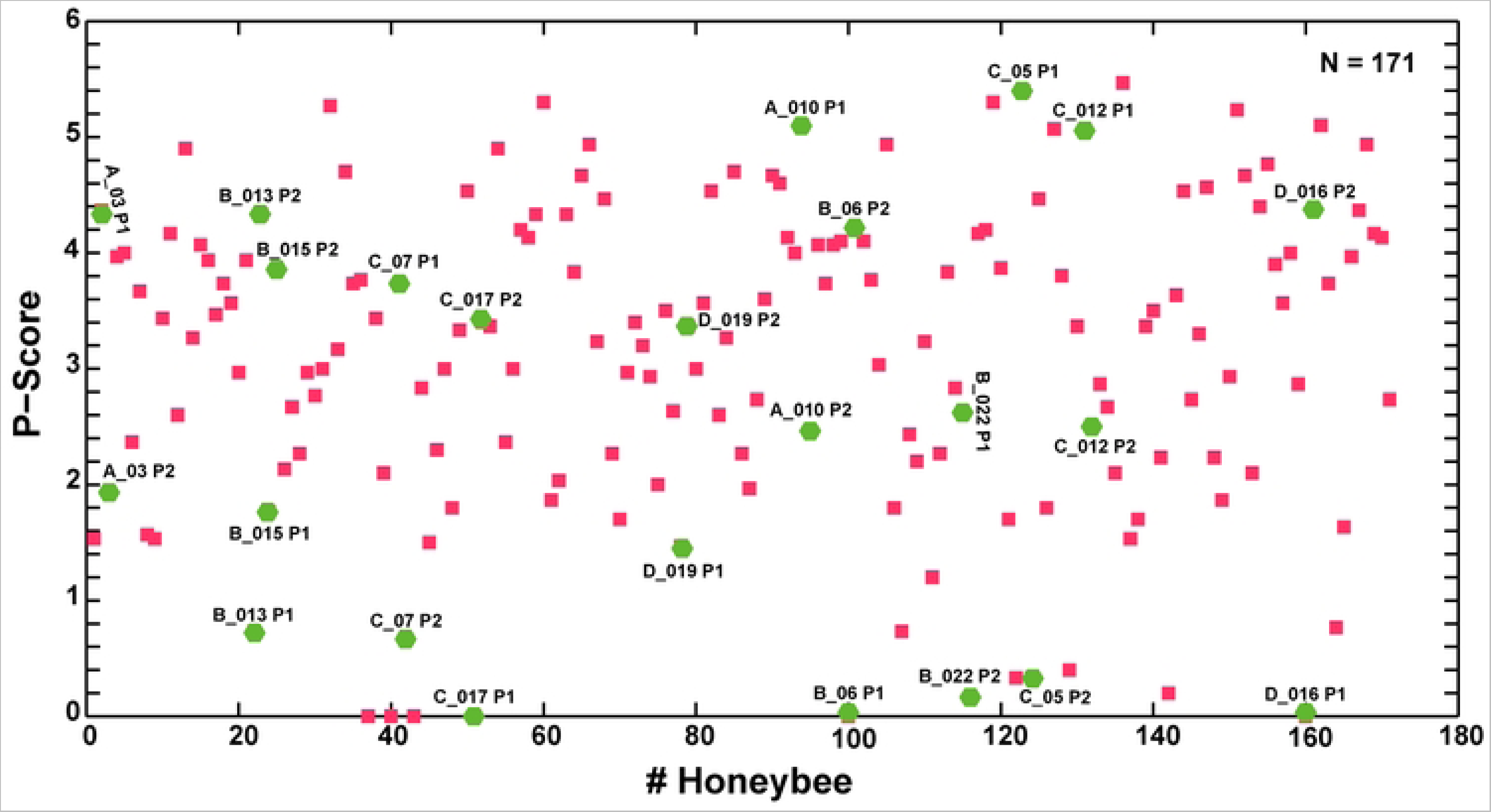
A sample of 24 bees (green hexagons with bee-identification numbers) out of the 171 bees (red squares) were selected for the genetic analysis. Among these 24 bees, 12 are high performers with P-scores ranging between 2.6 and 5.4 and the other 12 are low performers with P-scores ranging between 0 and 2.5.

**Table 6:**
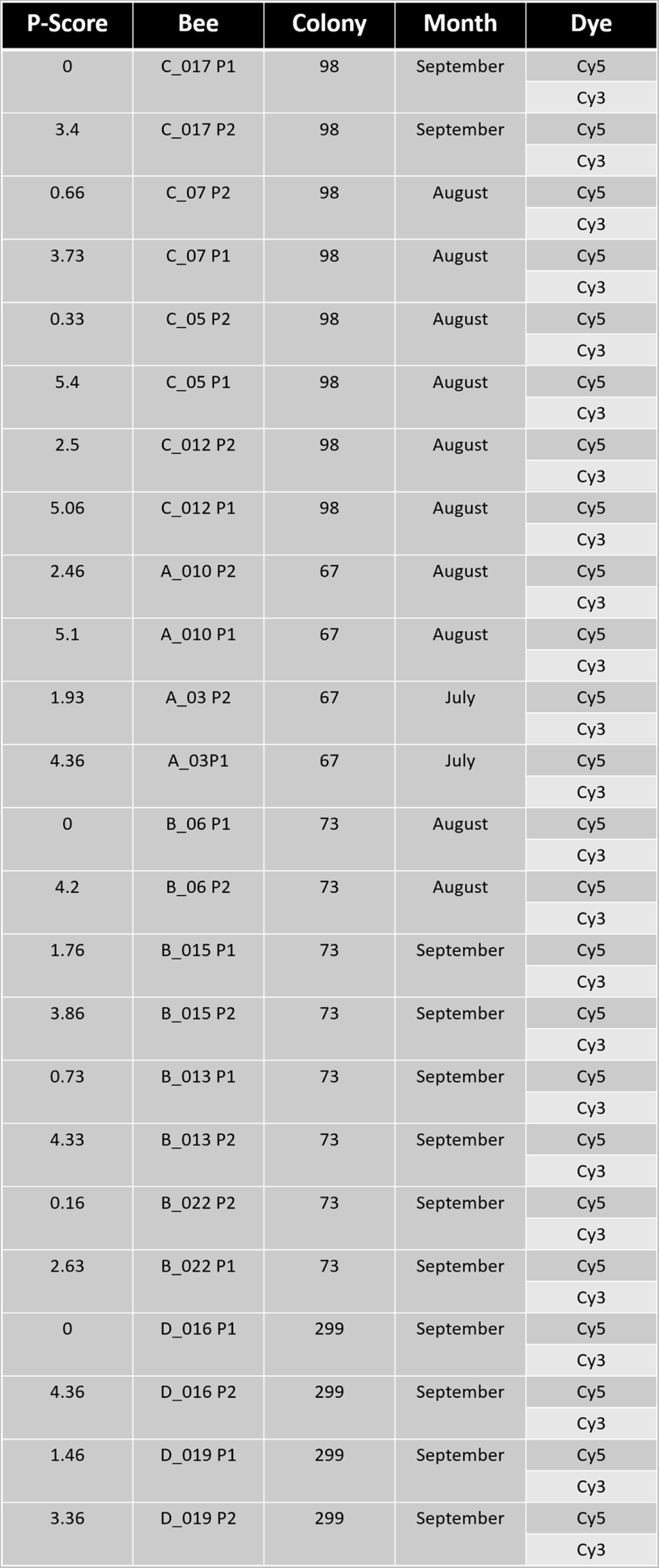
Selection of the pairs of bees from the four colonies, conditioned together (on the same day) in the different months of the experimental year, for the behavioral analysis based on their high and low P-scores. The column ‘Bee’ shows the identification number of the bees.

The microarray analysis of the behavioral design generated a list of 687 genes and the analysis of colony specific differences (genotype design) generated a list of 265 candidates. The behavioral list was sorted by removing 32 gene candidates also present in the genotype list, reducing the behavioral list to 652 candidates that were differentially expressed between the high and low performer bees with adjusted p-level varying between 0.0025 and 2.9×10^-7^ (Table S5). Most of the identified genes were characterized by slight differences in expression levels. The expression of 371 genes was reduced in the high performer bees, varying between 0.75× and 0.95× and the expression of 281 genes was increased between 1.05× and 2.46×. A list of 45 identified genes, including the candidates with the highest changes in expression levels, is presented in Table 7. They are potentially encoding proteins implicated in diverse biological functions, like neurotransmitter receptors for dopamine, tyramine, GABA and acetylcholine, proteins implicated in the synaptic release of neurotransmitters (VAMP, syntaxin, neuroligin) and amino-acid transporters (GABA, excitatory amino-acid transporters). Several candidates are implicated in cellular processes important for memory formation (proteasome complex, TOR pathway, protein translation) and in development (the transcription factor *mblk-1*, *minibrain*). Other interesting candidates like calmodulin, kinases, phosphatases are also known to be implicated in messenger pathways important for memory formation. Interestingly, genes which are regulated by growth hormone or juvenile hormone and malvolio, a gene implicated in the division of labor, also showed increased expressions in the high performer bees. Genes known to be absent from the mushroom body were also found in the list. The blue opsin gene expressed in the visual system probably originated from a contamination with the ocellar tract. Alpha-amylase, major royal jelly protein 5 (mrjp5) and glucose oxidase were the genes that showed the highest changes in expression; their expression levels were doubled in the high performer bees. These genes probably originated from a contamination with the hypopharyngeal glands, adjacent to the MB.

**Table 7:**
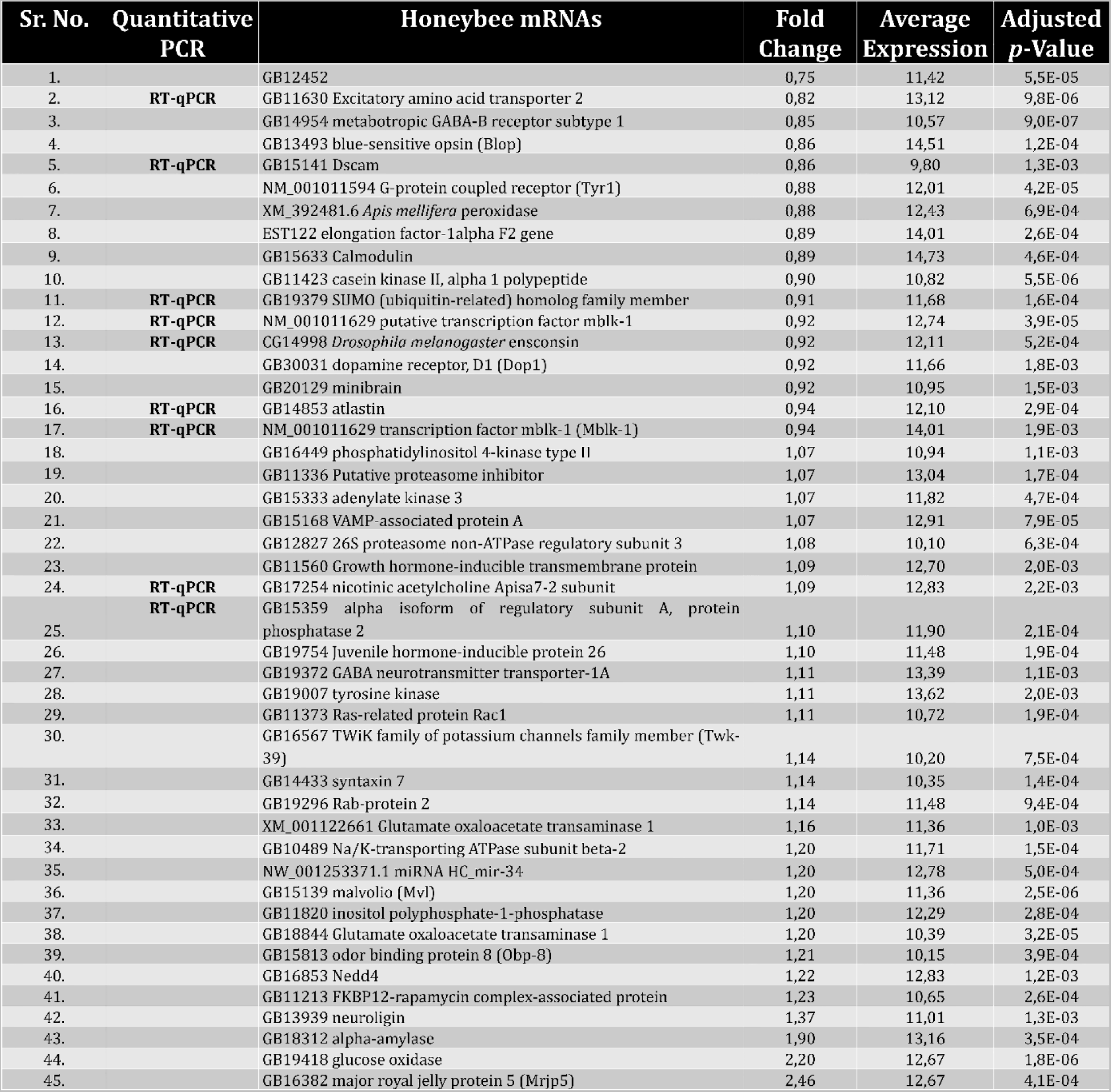
A selection of 45-identified genes from the sorted behavioral list that includes candidates with the highest changes in expression levels between the high and low performers. The table is showing the accession numbers and identity of the mRNAs deduced from the genome annotation (Honeybee mRNAs), the change in expression levels between the low and high performers (Fold Change), the averaged expression levels measured on the microarray, the adjusted *p*-values denoting significant changes in expression levels between the low and high performers, calculated from the microarray analysis, and the gene candidates that were tested by RT-qPCR (Quantitative PCR, RT).

Some genes of the whole genome honeybee microarray are linked to the orthologues of the *Drosophila* Flybase. In this manner, a list of 418 Flybase entries were generated from the sorted honeybee behavioral list (Table S7). The list was submitted to the KEGG Automatic Annotation Server (KAAS). The analysis revealed 600 enriched terms with terms related to sugar metabolism (e.g. pentose, fructose, galactose and sucrose), lipids metabolism (e.g. sphingolipid, arachidonic acid, linoleic acid), amino acid and nucleic acid metabolism (e.g. spliceosome, RNA transport, Ribosome) (Table S7). The list highlighted terms directly related to classical neurotransmitter systems (GABA, acetylcholine, glutamate, dopamine serotonin), olfactory transduction, axon guidance. Synaptic transmission (e.g. SNARE, Tight junction, GAP junction), the proteasome complex, several signaling pathways (e.g. Ras, MAPK, Ca^2+^, cAMP, cGMP, mTor). Many terms related to hormonal control (e.g. insulin, oxytocin, prolactin, adrenergic) and food processing (salivary-, gastric acid-, pancreatic-secretion, etc.) were also described. Several terms not directly related to the neuronal metabolic pathways underlying memory processes were also highlighted (e.g. immune response, longevity).

The behavioral gene list was also submitted to DAVID and the functional annotation chart revealed 76 statistically significant terms from the gene list. Some of them are related to microtubule associated complex, mitochondria, neurogenesis, plasma membrane, splicing, GTP activity, oxidoreduction and nucleic acid metabolism (Table S8). The eleven first clusters presenting enrichment scores between 1.59 and 0.81 highlight terms related to small GTP-binding protein domain, mitochondrion, PDZ domain, glycoside hydrolase, α/β hydrolase, spliceosome, protein phosphatase, actin cytoskeleton, neuroblast, small GTPase superfamily and lipid metabolism (Table S8).

### Validation of gene candidates by RT-PCR

Two low and two high performer bees from each of the four colonies (Table S9) were selected for the validation by real time PCR (RT-qPCR) of nine selected genes (Table 7). Low performers had scores varying between 0 and 3.33 and high performers between 3.8 and 5.4. and they were collected from July to October. None of the tested genes presented significant differences between the high and the low performer groups (Student t-test, p>0.05) (Figure S1).

## Discussion

A deeper understanding of the variations in individual behavior within species is attracting more attention owing to their significant contributions to the ecological traits of the species and their evolution [65] [66]. Heterogeneity in behavioral types or personality, which is related to the difference in cognitive ability of individuals, is a heritable feature in animals that controls important life-history traits, such as food intake, detecting prey, predators and mates, labor division, ability to respond better to the environmental change, and overall fitness [67] [68] [69] [70] [66]. Honeybees show individual variability in behavior, cognitive task specialization and consistency in learning proficiency across learning paradigms and stimulus modality ([71] [72] [73] [74] [75] and hence is an appropriate model system to study the nature of heterogeneity in cognitive abilities. In honeybee, it is already established that the group-averaged analysis of the learning data impedes the visualization of this heterogeneity in individual’s learning, viz. the learning rate, final level of learning, and the strength of retained memory [24] [25]. Response latency has been described to have the potential to underlie the heterogeneity in learning dynamics of the individual bees, which was explained by a heterogeneous Rescola-Wagner model [24] [25]. However, these analyses could not capture a clear picture on the identity of a salient behavioral feature(s) that determines the individualistic variability in learning ability in the honeybee populations. Here, we addressed this issue through a complex form of olfactory learning protocol and analyzed the data to understand the contributions of different learning-related features to the overall performance levels as well as investigated the correlations between learning performance and the expression patterns of learning-related genes in the brains of the individual honeybees.

### Methodological consideration

We trained the honeybees in a sequential olfactory conditioning protocol in which the animals received two consecutive stages of differential training with two different pairs of odors and each conditioning phase was followed by two retention tests in which we applied several dilutions of the rewarded (CS+) and unrewarded (CS-) odors. The protocol was specifically designed to test the responses of the individual bees rigorously for a longer period to find out the strength in their learned responses. The tasks of discrimination between the CS+ and CS- odors during the two stages of conditioning were relatively easier compared to the four sets of retention tests because of the use of pure odors and not their dilutions during the conditioning; thus the ‘Aha’ effect of learning led to clear separation in responses of the bees to the odors [76] [77]. Individual honeybees however had to perform consistently well throughout the procedure to reach a higher cumulative performance score (P-score) or they failed at different time points to score lower. This way we were able to distinguish between the low and high levels of olfactory learning performers present in the natural populations. Bees from all four colonies have learned the odor stimuli during the conditioning and discriminated well in the retention tests, however the discrimination task was more difficult when the cuticular volatile fatty acids (OA and LA) were used compared to the brood volatiles (OM and PEA). We speculate that learning of the differential contingencies of these two odors are difficult in the perceptual space of the honeybees owing to the structural similarities of these two compounds (both are 18-carbon, unsaturated essential fatty acids) [78]. This could be a prime reason for the bees to generalize more between the odors during the second half of the procedure in addition to the possible involvements of the non-associative components, like sensitization and hunger due to prolong testing of the animals for more than 6 hours with repeated sucrose stimulations. The associative components together with the non-associative counterparts make our procedure more stringent and specific in discriminating between the low and high levels of learning performances. We quantified the speed of learning of the CS+ odors and the levels of odor discriminability both during the two sets of conditioning and four sets of retention tests and investigated whether there is a single salient feature that persistently contributes to the overall learning performance of the individual bees and thus can be considered as an important feature that controls the heterogeneity in learning dynamics among the individuals of the natural populations. There are other complex olfactory learning protocols that were previously applied on the honeybees [79] [55] but we didn’t use them because either these protocols were not recording the bee’s responses for a longer period or they incorporated extinction components in learning, both of which were avoided. Though one may still argue that an extinction-learning component prevailed in our sequential conditioning design as we performed two consecutive sets of retention tests after each of the two differential conditionings. Indeed, bees responded less to the highest dilution of the CS+ odors (10^-3^) in the second compared to the first set of retention test (in the paired-retention tests), however no difference in memory recall was found for the training concentrations of the CS+ odors in the two consecutive sets of retention tests. Thus, we conclude that no extinction learning took place in our experimental design, which offered no time gap between the consecutive retention tests.

### Speed of learning explains the heterogeneity in learning and memory performance of the individual honeybees

A very few studies have systematically investigated the nature of heterogeneity or individualistic variation in the learning ability of honeybees. Tait and colleagues have reported individualistic variation in honeybee in the domains of olfactory and navigational learning, and maneuverability performance, however the authors did not conclude anything on the source of variation in the learning behavior of individuals [17]. Our analysis clearly demonstrates a large heterogeneity in the learning ability of the individual honeybees, which is structured with a continuous range of performance. Further, the variation in the speed of odor learning appears as the major component for this heterogeneity. The distribution deviate from an exact normal distribution. It demands further investigations if it is due to low sample size. Interestingly, normal distribution was reported for several cognitive capacities in human populations [80]. The distribution of P-score might reflect the complex social colony structure in honeybees where individuals at different developmental stages are involved in different tasks, which might require different degrees of specialization for particular learning modalities [72] [81] [82] . Indeed, variation among the individuals can emanate from the development of specific experience-dependent biases and thus, differences in physiology to detect, learn and respond to particular external stimuli [83]. Thus, in our experimental population of honeybees, which is made up of old and young foragers, including bees who were making transition from nursing to foraging (discussed later in this section), we found a range of performers for the olfactory learning and memory tasks.

The large heterogeneity in the learning dynamics of individual honeybees as seen in the previously generated learning data sets [24] [25] and in ours can probably be attributed to several factors, such as stimulus sensitivity, speed of learning, and memory retention capacity. We found that high performers responded more to the air puff stimulations without odor (filter paper only or with paraffin oil) than the low performers (Fig. 5). This result together with higher responses of the high performers than the low ones to the highest CS+ dilution (10^-3^) during the retention tests indicate possible higher olfactory sensitivity among the high performers.

The feature that has been found to mostly correlate with the overall performance was a higher acquisition rate during the first phase of conditioning. The speed of learning of the CS+ odor during the first differential conditioning (Acq-1) best predicted the cumulative performance of the individuals among all six quantified features. Individuals with higher speed of CS+ odor have not only discriminated the odors well during the first phase of differential conditioning, they also have higher retention scores and higher odor sensitivity. Although the honeybees also perform successfully in the second differential conditioning phase, only the speed of acquisition Acq-2 is strongly correlated to the speed of acquisition during the first differential phase. The discrimination index during the second acquisition phase (DisCond-2) and the discrimination during the second retention test (DisT-3,-4) were the least correlated to Acq1. These parameters are more influenced by the capability of an individual to adapt its behavior to new conditions and it adds an independent value to the selection of high performers.

Fast and slow learning are elaborately discussed in behavioral ecology in the context of behaviors that involve speed-accuracy trade-off. A fast learner can take more risk to gain more only for a short-term and even perhaps lacks accuracy, whereas a slow learner has better accuracy and it can perform the duty more safely with the notion for a long-term gain [84] [65]. The advantage of having slow learners has also been reported in a study on bumblebees where fast learning of visual information was found to be associated with lower number of days of foraging compared to the slow learning bumblebees, which described the fact that superior cognitive abilities may not be beneficial to build the reserve of the colonies [85].

We were unable to test the change over time of conditioned responses shortly after learning [86]. However, our results clearly show that bees, which started responding early to the CS+ odor compared to the late responders during the first conditioning, showed stronger and more specific odor memories in the short-term recall tests. The findings on memory specificity are contradictory to the findings of Pamir and colleagues [25] who used an absolute training procedure and found no difference in the specificity of odor memories between the early and late responders. We assume that this difference in results could be due to the differences in our conditioning methods and odor stimuli used, because the present sampling method is more controlled. We thus surmise from our results that the mechanisms underlying the speed of learning of odor stimuli are correlated to those controlling their discrimination and consolidations of memories.

### Genetic analysis in the mushroom body area of the high and low performers

An analysis of differences in gene expression in the mushroom body (MB) region of the low and high performers was performed holistically using microarray. The analysis identified 652 differentially expressed genes; most of them were characterized by small differences in expression levels between the high and low performer bees. This indicates that many genes each with small amplitude effects contribute to the high performer phenotype. We selected nine genes from the transcript list originally generated by the microarray method to further verify their expression by RT-qPCR. The presence of all nine transcripts could be confirmed by RT-qPCR, but no significant expression changes were detected in the high and low performer groups by this technique (Figure S1). These results illustrate that behavioral traits are influenced by small changes in gene expression of multiple genes [84]. This particularity is also a feature described for neurological disorders [87] [88].

The KEGG and DAVIDD analyses revealed similar terms that were in accordance with identified genes from the gene list. Nervous system specific terms are related to plasma membrane and synaptic transmission, illustrated by neurotransmitter receptors (GABA, dopamine, octopamine, acetylcholine, glycine and serotonin) and to synaptic components related to neurotransmitter release (e.g. SNAREs). Terms related to LTP, LTD, and neurogenesis were also revealed, as well as terms related to olfaction, taste and photo-transmission. The latter might indicate contamination with tissues adjacent to the MB. Indeed, some of the terms can also be related to increased metabolic activity, like sugar metabolism, or to mitochondria and cellular respiration. The analysis of the gene list also supports that the MB dissections were contaminated with hypopharyngeal glands. We identified higher levels of the major royal jelly protein 5 gene (*mrjp5*) in the high performers. However, However, all *mrjp*s are produced in the hypopharyngeal glands except for *mrjp3* that is produced in the Kenyon cells of the MB of nurse bees [89] [90] [91]. Alpha-amylase and glucose oxidase are produced in the hypopharyngeal glands and are also overexpressed in the high performers [92] [93] [91]. Several intracellular signaling cascades, known to be implicated in learning and memory, were also highlighted like several second messenger pathways (TOR signaling and the ubiquitin proteasome complex). These pathways regulate, among others, translation- and transcription-dependent processes that are important in memory processes and are highlighted in the different lists. However, these processes are not restricted to neurons and might thus highlight activities in adjacent tissues.

The hypopharyngeal glands are important for food processing in honeybees [94] [95] [96, 97]. The genes *mrjp*1-4 and *7* are more active in the nurse bees compared to the foragers [90]; [91]. In foragers, α-amylase and glucose oxidase are associated with the conversion of nectar to honey [92] [93]. The change in expression of these enzymes is dependent on the age and the task. Interestingly, *malvolio*, a protein influencing labor division in honeybees, is also overexpressed in the high performers [98] [99]. This manganese transporter was discovered in *Drosophila* and is known to influence sucrose responsiveness [100]. Thus, we hypothesize that the high performer bees are in the transition from nursing to foraging activities, which is supported by the highlighted genes and terms related to hormonal pathways. It is also known that immunity changes during this transition [93] [101] and in terms related to immune response were highlighted in the gene ontology analysis.

In a previous study, Whitfield [57] and colleagues analyzed behavioral maturation in the honeybee by comparing the gene expression profiles in the brains of bees in different settings: nurses and foragers of specific ages compared in crossed conditions as well as nurses treated with different chemicals, described for their roles in the onset of foraging, Methoprene (an analogue of juvenile hormone), manganese and cGMP. Both settings were reported to associate with the molecular pathways of *malvolio* and *foraging*, however cAMP was found to play no role with the onset of foraging [57]. We compared our gene list to the gene lists of them. The comparison of gene sets characterized by a significance level < 0.05 did not reveal any significant overlap. By restricting the gene sets to candidates, characterized by a significance level < 0.001, significant effects were described (Table 8). A significant correlation was found for Methoprene treatment, a condition also implicated in the transition from nursing to foraging. However, there was no significant overlap with manganese or cGMP treatment neither with the comparison between 17 days old foragers and nurses (d17F/d17), which are conditions specific for foragers. Significant overlap was found with gene sets specific for juveniles, which are still active with hive duties but able to perform the transition from nursing to foraging (d4/d8 and d12/d17). This is a hint that high performers belong to a group of young bees that were in the transition to foraging activities. The animals were caught at the end of the afternoon, when mostly foragers perform outbound flights. Thus these bees might correspond to young individuals beginning their foraging activity. It is known that young foragers and precocious forager honeybees learn olfactory stimuli better than the nurse bees [102] [72]. However, less is known about differences between young and old foragers. Only a few studies addressed this question [103] [104]. The correlation with the gene set specific for the cAMP second messenger pathway can explain the higher sensitivity of the high performer bees compared to their low performing counterparts. Indeed, this second messenger pathway is associated to modulatory neurotransmission mediated by the biogenic amines, like serotonin, octopamine, dopamine, and tyramine that are implicated in arousal, sensitivity, and learning performances [105] [72] [106]. The cAMP second messenger pathway is linked to some of these neurotransmitter systems, which might explain why the high performer bees are responding more to olfactory stimulation with filter paper and paraffin oil. Although, the selection of high and low performers was biased for olfactory sensitivity, the differential conditioning procedure demonstrates that learning and memory are specific. The analysis of the gene expression profiles of high performers suggests that they are young individuals at the transition to foraging activities. The brains of the high performers present differences in several neurotransmitter systems, components involved in synaptic transmission and in signaling pathways important for learning and memory. Further investigations are needed to describe more precisely the differences between young and old foragers.

**Table 8:** Comparison of the gene list with adjusted *p* value < 0.001, described in this study (n = 326) and in the study of Whitfield and colleagues (2006). The number of differentially expressed genes identified for specific conditions are presented (Total differentially expressed genes). These were treatments with Methoprene, Manganese, cGMP, cAMP, and cAMP vs cGMP. Age effects were also considered in the nurse bees of different ages (days, d). Seventeen days old nurses and foragers were also compared (d17F/d17). We evaluated if the number of genes overlapping between the sets identified in this study and those of Whitfield (Gene overlap) was random with a χ^2^ test. Significant differences have a *p* value < 0.05.

| Treatment | Methoprene | Manganese | cGMP | cAMP | cAMP/cGMP |
| --- | --- | --- | --- | --- | --- |
| Total Differentially Expressed Genes | 481 | 509 | 911 | 129 | 327 |
| Gene Overlap | 24 | 11 | 28 | 7 | 10 |
| $\chi^2$ | 13.8566 | 0.1564 | 1.7335 | 4.9555 | 0.5665 |
| *p*-Value | 0.000197 | 0.692517 | 0.187961 | 0.026008 | 0.451641 |
| Age | d1/d4 | d4/d8 | d8/d12 | d12/d17 |  |
| Total Differentially Expressed Genes | 2613 | 1260 | 342 | 160 |  |
| Gene Overlap | 69 | 44 | 11 | 8 |  |
| $\chi^2$ | 0.6338 | 6.6813 | 0.9272 | 4.5344 | |
| *p*-Value | 0.425968 | 0.009743 | 0.33559 | 0.03322 |  |
| Age | d17/d1 | [d8,d12,d17]/d1 | d17F/[d8,d12,d17] | d17F/D1<br>7 |  |
| Total Differentially Expressed Genes | 3745 | 3563 | 3575 | 2414 |  |
| Gene Overlap | 101 | 97 | 94 | 62 |  |
| $\chi^2$ | 1.6149 | 1.8049 | 0.8545 | .2534 | |
| *p*-Value | 0.203798 | 0.179123 | 0.355274 | 0.614713 |  |

## Acknowledgements

We are grateful for the collaboration with Dr. Evren Pamir in the course of analyzing our data. We also thank Mr. Peter Knoll for the help with the bees. We acknowledge the collaboration with Prof. Kaspar Bienefeld, Mr. Fred Zautke, and Dr. Caspar Schoening from the Länderinstitut für Bienenkunde Hohen Neuendorf e. V. for creating the beelines and delivering them to our institute. We are grateful for the collaboration with Dr. Evren Pamir in the course of analyzing our data. We also thank Mr Peter Knoll for the help with the bees.

NKC was supported by the German Federal Ministry for Education and Research (BMBF) within the framework of the FUGATO program (ref. no. 0315124A), GL by the grant Deutsche Forschungsgemeinschaft LE 1809/2-1 and RM by the grant Deutsche Forschungsgemeinschaft Me 365/41 and the Freie Universität Berlin.

